# TissueFormer: Extending single-cell foundation models to predict population-level phenotypes

**DOI:** 10.1101/2025.08.17.670735

**Authors:** Ari S. Benjamin, Anthony Zador

**Affiliations:** Cold Spring Harbor Laboratory, Cold Spring Harbor, NY 11724, United States

**Keywords:** spatial transcriptomics, single-cell foundation models, sample-level diagnostics, brain annotation, tissue labeling, AI for biology

## Abstract

**Background:** Single-cell RNA sequencing technologies have enabled unprecedented insights into gene expression and opened new pathways for diagnostics and tissue annotation. At present, most computational approaches for interpreting single-cell data predict labels or properties based on isolated single-cell transcriptomic profiles. This approach overlooks the cellular composition within a sample, which is often critical for inferring tissue identity or other sample-level phenotypes.

**Results:** To address this limitation, we introduce TissueFormer, a Transformer-based neural network that infers population-level labels from groups of single-cell RNA profiles while retaining single-cell resolution. We applied TissueFormer to two tasks: predicting COVID-19 severity from single-cell RNA sequencing of blood samples, and predicting cortical area identity from spatial transcriptomic data in mouse brains. TissueFormer outperformed single-cell foundation models and machine learning methods applied to pseudobulk and cell type composition.

**Conclusions:** TissueFormer’s higher performance promises more accurate diagnostics and enables the automated construction of high-resolution brain region maps in individual mice directly from spatial transcriptomic data. Applied to mice with developmental perturbations to visual input, these maps revealed a significant reduction in predicted visual cortex area, illustrating how individual differences in neuroanatomy can be quantified. More broadly, TissueFormer provides a framework for predicting any population-level phenotypes which are influenced by cellular diversity and tissue-level organization.

## Background

Many important features such as tissue identity, disease state, or treatment response are defined at the level of cell populations but are driven by patterns of single-cell phenotypes. Observations of population-level phenotypes can thus be used for the annotation and classification of tissue samples. For example, the proportion of cell types in a population is a useful biomarker for clinical features such as inflammation (***Templeton et al., 2014***). In the brain, an important problem in this class is the annotation of brain areas which differ in cell type composition (***Yao et al., 2021b***). However, readouts depending on cell composition currently remain limited to a handful of canonical patterns identified through decades of observation rather than discovered systematically through large-scale, data-driven analyses.

With the increasing availability of single-cell RNA sequencing, it is now possible to measure patterns in single-cell properties at high resolution across large numbers of cells per sample (***Svensson et al., 2018***). This rich granularity presents a tradeoff for the analysis of phenotypes across tissues, samples, or individuals, each of which contains many sequenced cells. At one extreme, many computational pipelines focus on single cells in isolation, taking a cell profile as input and outputting a label specific to that cell. This is the approach taken by most recent single-cell ‘foundation models’ of mRNA transcription data (***Lopez et al., 2018; Connell et al., 2022; Cui et al., 2023; Theodoris et al., 2023; Rosen et al., 2023; Yuan et al., 2024; Schaar et al., 2024; Ito et al., 2025; Hsieh et al., 2024***), reviewed in (***Szałata et al., 2024***). These models are fundamentally single-cell in design, and as a result they are unable to detect signals that emerge only from the *composition* of cells present.

An opposing strategy is to first compute summary statistics about cell populations across tissues or samples, then use these for downstream analysis and classification. One such statistic is the average expression, or pseudobulk expression profile, as is frequently used in differential state analysis (***Crowell et al., 2020***). Alternatively, one may calculate compositional features about a population, such as cell type frequencies, ratios of cell types, putative cell-cell communication networks, and cell type specific pathway scores (***Cao et al., 2022***). While informative, both pseudobulk and cell composition measures obscure single-cell granularity and rely on the researcher to specify which compressed representation of a population of cells is likely to be informative about sample labels.

In order to maximize the potential of single-cell data, it is important that methods combine the transfer-learning capabilities and end-to-end trainability of single-cell foundation models with the ability to compare cells across a population. Certain models exist which are end-to-end trainable and can compare across cells, such as CellCnn (***Arvaniti and Claassen, 2017***), scAGG (***Verlaan et al., 2025***), and ScRAT (***Mao et al., 2024***), but these models are not compatible with existing foundation models. The ability to incorporate knowledge from pretrained models is especially important given the high dimensionality of the inputs when predicting sample-level phenotypes. In each independent sample, the input dimensionality is on the order of the number of cells per sample times the number of genes per cell. By first pretraining on self-supervised tasks, foundation models effectively reduce this dimensionality by establishing a learned prior over the representation of each cell.

To bridge these approaches we present TissueFormer, a neural network architecture that analyzes groups of single-cell RNA profiles collectively while utilizing knowledge embedded in pre-trained single-cell foundation models. The model incorporates a module based on the Geneformer architecture for single-cell analysis (***Theodoris et al., 2023***), which can be initialized with pre-trained weights. During processing, each cell is given a learnable representation with this pretrained module. These cell representations are then processed by several further layers of self-attention, allowing the model to learn to attend to the relevant cells and their interactions. Importantly, the output of Tissueformer is invariant to the order of cells presented as input. These design choices allow TissueFormer to learn arbitrary functions from observations of many cells to sample-level phenotypes while simultaneously leveraging pretrained single-cell foundation models.

To validate our approach, we applied TissueFormer to two prediction tasks in which cellular diversity is likely to be crucial. In the first task, we link spatial transcriptomic data to Allen Brain Atlas labels of regions of the mouse cortex. This task is a competitive benchmark for the supervised classification of sampled tissues, though it is important to note that clustering or alignment-style methods have also been developed for tissue annotation (e.g. ***Biancalani et al. (2021); Hu et al. (2021); Lee et al. (2025)***). As a second task, we compiled three datasets of single-cell transcription within blood samples of patients with COVID-19 infections of varying severity, and sought to predict infection severity from gene expression (***COMBAT, 2022; Ren et al., 2021; Stephenson et al., 2021***). On both tasks we found that TissueFormer outperforms both standard supervised methods as well as fine-tuned single-cell foundation models at predicting sample-level annotations. These results highlight the importance of sample- or tissue-level features in transcriptomic analysis and establish a framework for integrating single-cell foundation models into higher-order biological investigations. When applied to brains of mice with developmental perturbations, TissueFormer’s predicted cortical maps further revealed individual differences in neuroanatomy that are consistent with past analyses of compositional shifts in these brains (***Chen et al., 2024***).

## Methods

Our primary goal is to design and evaluate a model which is generally applicable to the supervised problem of sample or tissue annotation from single-cell data. In what follows, we first describe the particular characteristics of this problem that make it challenging for standard methods.

### Multi-cell supervised problem setting

A typical problem in single cell analysis is relating mRNA expression levels in each cell to a cellular property such as cell type identity (***Figure 1***a). If we denote the cell-by-transcript matrix of counts as *C*, where each row of *C*_*i*_ contains the mRNA transcript counts for a single cell, this problem seeks to find a function *f* that maps each row *C*_*i*_ (a cell) to a particular value *y*_*i*_ corresponding to a label. For example, *y*_*i*_ could represent the cortical area from which a given cell was obtained.

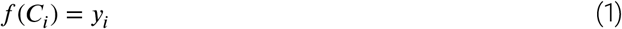

**Figure 1.**
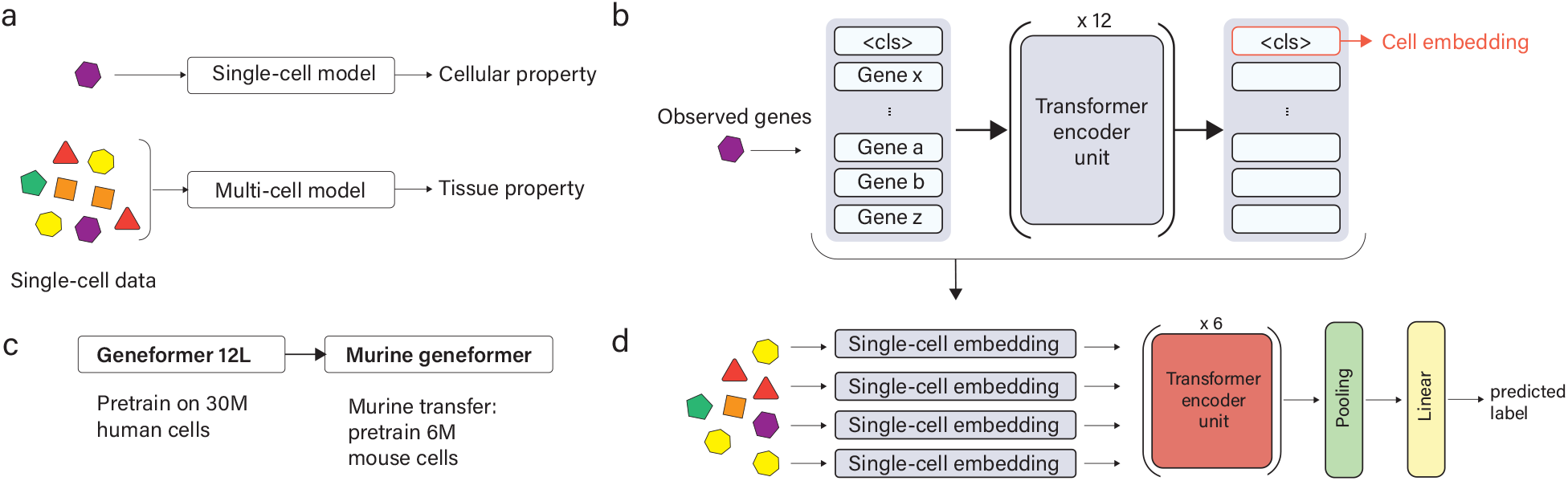
TissueFormer is a Transformer-based neural network to analyze groups of single-cell RNA profiles. a) Single-cell foundation models focus on cellular targets, yet many properties of tissues relate to cellular diversity. b) TissueFormer learns to extract critical information from each cell into a cell embedding vector. The module that extracts these is in architecture identical to Geneformer, a single-cell foundation model. First, cells are processed by a normalized rank-ordering of their genes into a ‘sentence’ of gene tokens, prefixed by a ‘<cls>’ token to pool gene information, then processed by several transformer layers. c) Experiments in this manuscript use a pretrained single-cell model we call Murine Geneformer, created by adapting the 12-layer Geneformer model (pretrained on human cells) to mouse cells via pairing orthologous genes and further pretraining. d) TissueFormer architecture. Cell embeddings for all cells in a group are fed to 6 Transformer encoding units before average-pooling and a linear readout.

In some problem settings, it is advantageous to use information from many cells. In this multi-cell setting, the characteristic equation relates sets of cells to labels. For spatial transcriptomic data, for example, a set might refer to a group of cells within some spatial distance of one another. Denoting *G*_*i*_ as the *i*th such group, and {*c* ∈ *G*_*i*_} as the set of cells in this group, the classification or regression problem seeks to find a function *f* that maps each set of cells to its label *y*_*i*_.

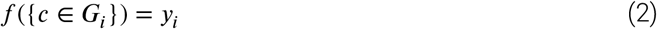

The fundamental difficulty of the multi-cell problem is its high dimensionality. Relative to the single-cell problem, the dimensionality of input examples increases from the number of genes *D* to *D* × |*G*|, where |*G*| is the number of cells in a group. Meanwhile, the number of independent labels decreases from the total number of cells to the number of disjoint groups, i.e. from *N* to 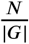. The ratio of input dimensions to independent examples, a proxy for the difficulty of the problem, thus increases as the group size squared, |*G*|^2^.

Most strategies for the multi-cell problem explicitly reducing its dimensionality through aggregation (as in pseudobulk analyses) or summary statistics (e.g. ***Cao et al. (2022)***). However, these strategies destroy information which could be critical in multi-cell problems. Pseudobulk methods obscure the variance in the signal, whereas cell type composition or other such signals are fixed and non-adjustable representations of single-cell profiles and are incapable of observing transcriptional dysregulation within cell types. To take full advantage of single-cell data, a model should ideally be able to attend to any aspect of the full RNA transcriptome in each cell while comparing across the population.

It is nevertheless crucial take steps to constrain the distribution of learned functions so as to circumvent the curse of dimensionality in multi-cell prediction problems. One way to address this is by pre-training a foundation model on a self-supervised task before downstream supervised learning. If this self-supervised task is sufficiently similar to the supervised task at hand, the pretrained model weights will be close in weight space to a good solution for the supervised task. This reduces the number of training steps and data needed in the supervised task.

Recent related work has introduced several foundation models tailored for spatial transcriptomic data. Because these models observe groups of cells, they can be seen as solutions to a subcase of the multi-cell problem in Equation 2. For example, scGPT-spatial is pretrained to predict gene expression in held-out cells in the local neighborhood (***Wang et al., 2025***). Unlike TissueFormer, scGPT-spatial averages the embedding of several cells in the local neighborhood for the inter-cell problem, rather than employing self-attention across cells, and thus cannot compare compositional signals. HEIST (***Madhu et al., 2025***) and CI-FM (***You et al., 2025***) define graph neural networks over a hierarchical graph spanning both nearby cells and their genes, which in principle enables comparative processing across cells. As explained below, TissueFormer uses Transformer layers and not graph neural networks, making it compatible with single-cell foundation models trained on traditional scRNA-seq data. Unlike these works, the design of TissueFormer is not restricted to spatial transcriptomic data and is applicable to any sample-level phenotype prediction problem.

### TissueFormer architecture

TissueFormer is a neural network architecture tailored for groups of single-cell RNA transcriptomic profiles (***Figure 1***). It is able to attend to information within each cell’s relative mRNA transcription profile, yet also look across cells to extract information from the diversity and frequency of mRNA profiles in the group. Architecturally, this is accomplished by first applying a ‘single-cell module’ to extract information from each cell, then attending across cells in further layers. This combination of intra- and inter-cell expressivity creates a regression problem of high dimensionality, yet this is mitigated by the bottleneck architecture and the easy loading of pre-trained single-cell foundation models into the cell embedding module.

The building block of TissueFormer which performs these capabilities is the Transformer block implementing self-attention (***Vaswani et al., 2017***). Compared to older neural network architectures like Multi-Layer Perceptrons (MLP), Transformers are distinguished by the shape of their inputs. Whereas an MLP observes vector-shaped inputs (i.e. a vector of mRNA transcript counts), Transformers operate on lists of vectors, or equivalently, on matrix-shaped inputs. Self-attention acts to select which vectors in the list are most relevant in the current context.

In order for single-cell data to be compatible with Transformers, each cell’s vector of mRNA transcript counts *c*_*i*_ must be expanded into a list of vectors. Multiple strategies for this conversion are possible for single-cell data, as reviewed in ***Szałata et al. (2024)***. Here we use a rank-ordering approach as in Geneformer (***Theodoris et al., 2023***). Each cell’s expressed genes are ordered by their relative expression normalized by their median expression in a pretraining corpus. Each gene name in this ordered list is then mapped to a high-dimensional embedding vector via a learnable dictionary. While alternatives to rank encoding have been tested in other foundation models, rank encoding is common in bioinformatic pipelines and has generally been found to be robust to common sources of experimental variability (***Ballouz et al., 2015***).

The single-cell module within TissueFormer is designed to compress each cell’s unique and relevant information into a *cell embedding* for later processing (***Figure 1***b). This is achieved by prefixing to each cell a special token named the <cls> token, following standard design choices (***Devlin et al., 2019***). The <cls> token instantiates a working space for the accumulation of information from all genes. During learning, any error-corrective information passed backwards is bottlenecked through the cell embedding, ensuring it collects the necessary information from each cell. In sum, the cell embedding vector which is passed to later layers can be written as a function of the mRNA transcript count vector *c*_*i*_ as:

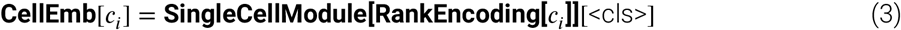

The single-cell module shares the Geneformer architecture, meaning that any trained single-cell model can be loaded into this module. This gives TissueFormer the ability to incorporate model weights and biological knowledge from pretrained single-cell foundation models. In our experiments we used a version of Geneformer adapted for mouse brain tissue. We first downloaded the Geneformer 12L model, which was trained to predict the identity of randomly masked genes (15% of total genes per cell) on a corpus of 30 million human cells. We then adapted this to mouse tissue by repeating this methodology on a hand-curated dataset of approximately 3.5 million mouse cells. Using the CellXGene Census explorer (***Program et al., 2025***), we aggregated data from several publications (Table 1) (***Yao et al., 2021b; Govek et al., 2022; Kozareva et al., 2021; Steuernagel et al., 2022; Yao et al., 2021a, 2023; Zhang et al., 2021***). After tokenizing (see below), we then trained the Geneformer architecture on this dataset with an identical masked gene prediction objective. Models were trained for 3 epochs using AdamW, a learning rate of 0.001, and a batch size of 32 cells on two Nvidia V100 cards. We tested training on the murine dataset from scratch as well as finetuning the human model. In order to finetune the human model, we gave mouse genes the same initial embedding vector as their human orthologs, where available, and gave unmatched genes random initializations. Orthologs were obtained from Ensembl (***Harrison et al., 2024***), with the first selected if multiple orthologous genes existed. This transfer from human genes significantly improved the model compared to training on mouse genes alone as measured with the cross-entropy loss on 100,000 held-out cells from the mouse brain datasets. We call this finetuned Geneformer pretrained model ‘Murine Geneformer’. When used within TissueFormer, Murine Geneformer serves as the single-cell module; at group size *N* = 1, TissueFormer reduces to Murine Geneformer with a classification head.

**Table 1.**
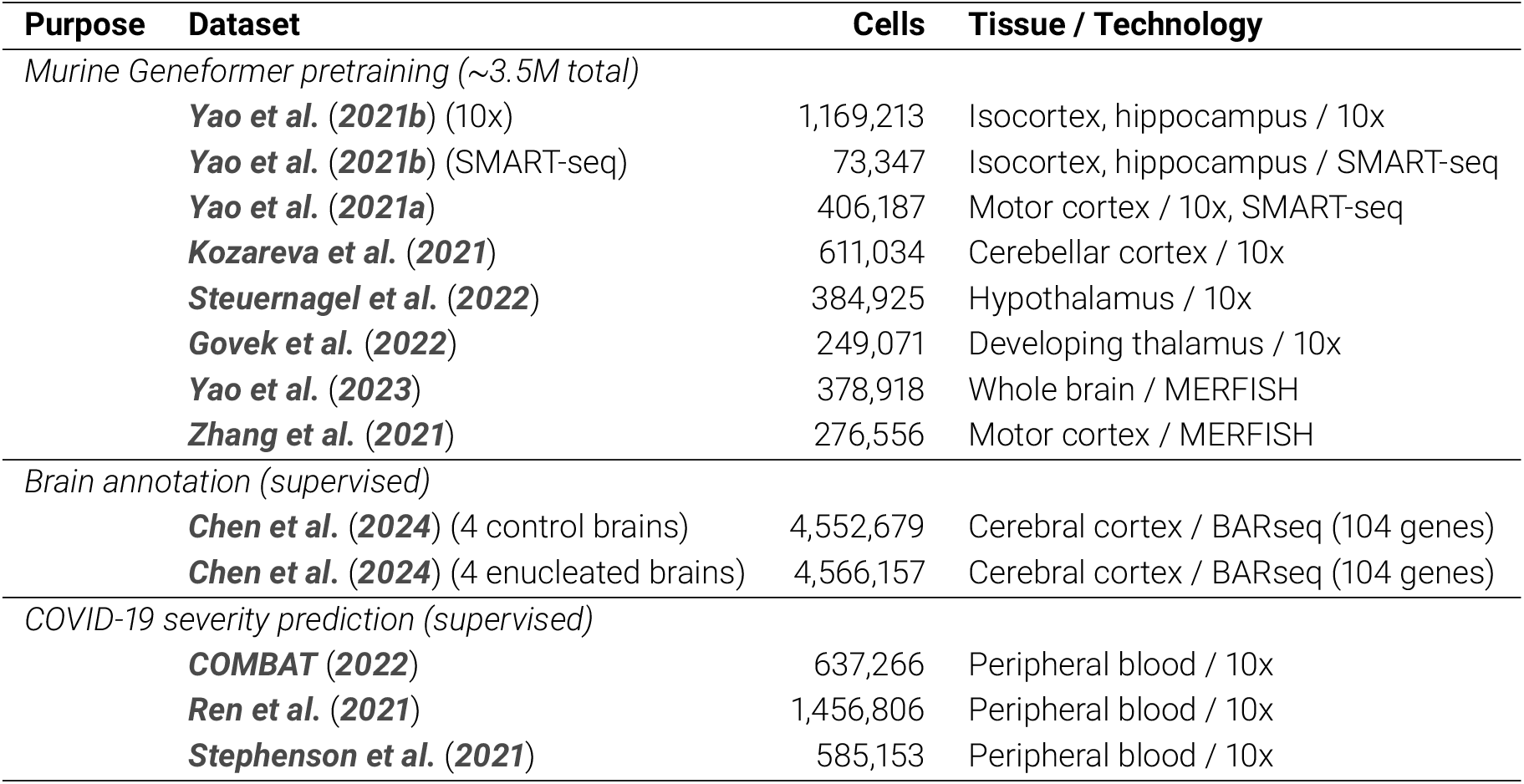
Summary of datasets used in this study.

TissueFormer attends to information across cells by processing each cell embedding with several additional Transformer layers (***Figure 1***d). At this stage, TissueFormer is invariant to the order of cells presented to the model, seeing them only as a bag or set of cells. The output of any Transformer is inherently equivariant with respect to the order of the input embedding vectors (i.e. rearranging the inputs causes the outputs to be rearranged in the same order). In order to attain invariance to cell order, such that rearrangements to the order have no effect on the output, the output of the Transformer layers is pooled by averaging over cells. A final step is a linear classification layer for classification problems. Collectively, these design choices allow TissueFormer to attend to any relevant information in each cell’s transcriptome in a context-dependent manner based on the diversity of cells present and the problem at hand.

Taken together, the equation representing TissueFormer can be summarized as:

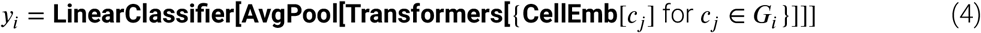

### TissueFormer architecture: hyperparameters of this study

TissueFormer contains two main components: a single-cell module and an aggregation module. In this sense it can be viewed as an end-to-end trainable hierarchical model above single-cell foundation models.

The single-cell module is based on the BERT architecture. We used identical hyperparameter settings as used by Geneformer 12L (***Theodoris et al., 2023***), allowing pretrained weights to be readily employed. Specifically, we used a 12-layer BERT network with ReLU activations, hidden layer sizes of 512, and a context window of up to 2048 genes. Each Transformer had multi-head attention with 8 heads, and the inputs are appended with sinusoidal positional embeddings.

The aggregation model pools across single-cell embeddings. The aggregation module had 6 Transformer layers, each with a hidden size of 768 units. All training runs on the classification task used the Adam optimizer with a learning rate of 0.001 and a linear learning rate decay following a learning rate warmup with warmup ratio 0.1. Our implementation relies on the Hugging Face Transformers library (***Wolf et al., 2020***), based on Pytorch (***Paszke et al., 2019***). We used Hydra for configuration management (***Yadan, 2019***).

### Application to cortical annotation

The mammalian brain is often partitioned into regions with distinct functional roles, connectivity, and cellular composition. In any particular brain, variations in neuroanatomy may arise due to genetic factors as well as due to the animal’s environment. These variations underlie the evolution of innate behavior and reflect the aptitude for adaptation within a lifetime.

Current methods for the annotation of single brains have certain well-known limitations. One common approach is to register a brain into the average-brain template of the Common Coordinate Framework (CCF), a 3D reference coordinate system constructed from over 1,000 reference mouse brains (***Wang et al., 2020***). Once transformed into CCF coordinates, one can visualize the area boundaries of the Allen Brain Atlas, which again reflect a canonical, average brain. The drawback of this approach is that it obscures individual variation, especially the relative size of brain areas to one another. An alternative approach which enables individual variability is to employ experimental methods that interrogate only a small brain region in each animal, for example, focal viral projection tracing (***Xu et al., 2020***) or two-photon functional imaging for a particular functional modality like visual responses (***Kalatsky and Stryker, 2003***). While accurate, these methods’ limited spatial scope prevents a holistic assessment of anatomy across the entire cortex.

An alternative strategy that balances whole-brain coverage with single-animal individuality is to perform *in situ* single-cell RNA sequencing across a single brain. This method modernizes a classic approach of defining neuroanatomy through differences in cell type composition, an idea that goes back at least to Brodmann’s brain atlas (***Brodmann, 1909; Zilles and Amunts, 2010***). Currently, the technical efficiency of spatial transcriptomics is now high enough to allow the profiling of cells across an entire mouse brain, and furthermore cheaply enough to cover multiple brains in single studies (***Chen et al., 2024***). Given such datasets, the remaining challenge in a robust pipeline for labeling brain areas is computational in nature.

To investigate whether TissueFormer could serve this purpose, we applied TissueFormer to predict the brain region of cells from their transcriptomes as characterized in a recent dataset of whole-brain spatial transcriptomics of the brains of several mice (***Chen et al., 2024***). Over 1 million cells across the brain, largely in the left hemisphere, were profiled in each of eight mice, four of which were raised in normal rearing conditions. Each cell profile contains counts from a panel of 104 genes selected for their ability to distinguish cell type clusters in excitatory cells. With multiple separate brains and whole-cortex coverage in one hemisphere, this dataset enables a test of whether individual differences in neuroanatomy can be resolved with spatial transcriptomics.

In addition to mRNA counts, this dataset also contains labels of the brain area of each cell as established through a registration into the Common Coordinate Framework (CCF). Each brain in ***Chen et al. (2024***) was previously registered to the CCF using standard software that defines a non-rigid smooth deformation from raw slice coordinates into the CCF. This step normalizes differences in gross anatomy and brain size. Importantly, once transformed into CCF coordinates, one may associate the topography of a new brain with the Allen Brain Atlas to obtain brain area annotations (***Wang et al., 2020***). These CCF annotations provide the labels we use for training.

It is important to note that CCF boundaries may not reflect ground truth neuroanatomy. While CCF registration accounts for overall brain shape, it does not account for more fine-grained differences such as the relative size of brain areas or a shift in a single boundary in one animal versus another. Nevertheless, these labels are sufficiently accurate to allow a comparison of computational methods. It is possible as well that the ‘errors’ of a model trained on such data will in fact represent a more accurate annotation than the CCF, a possibility we will return in ***Figure 3*** in which we examine the predictions of TissueFormer on held-out brains.

### Data and tokenization

All datasets used in this study are summarized in Table 1.

#### Data description: mouse brains

The data published in ***Chen et al. (2024)*** contain spatial transcriptomic data obtained via BARseq. BARseq is a probe-based *in site* sequencing technology based on Illumina chemistry. BARseq establishes mRNA transcript counts in cells based on the identification of probe sequences at spatially resolved locations, followed by cell segmentation and assignment of genes to cells. In this dataset, probes were selected to resolve 104 marker genes chosen to resolve excitatory cells in cortex. After quality control, the overall dataset contains 10.3 million cells across nine brains, including a pilot brain, four brains of mice raised in normal rearing conditions, and four brains of mice raised following monocular enucleation.

After downloading spatial transcriptomic data from ***Chen et al. (2024)***, we first converted all files into AnnData format (***Virshup et al., 2023***) with a custom script. We selected the four mice raised in control conditions, then selected only cortical cells, resulting in over 1.6 million cortical cells across 4 mice. Several metadata annotations for each cell were computed in the original publication. These include cell type annotations at 3 hierarchical levels, and cellular location within each brain slice in ‘slice coordinates’ as well as in CCF coordinates after each brain was registered to the Allen Brain Atlas. This dataset also contains the location within warped ‘flatmap’ coordinates, which use the butterfly projection from the `ccf-streamlines` python package to place cells in a 3D coordinate space in which cortical depth is the third axis (***Wang et al., 2020***).

For plots in which cells were colored by cell type (e.g. ***Figure 3***), we assigned to each cell type a color so that similar cell types would have similar colors. Specifically, the perceptual distance between any two cell types’ colors was chosen to match the correlation between the cell types’ pseudobulk expression vectors. Unlike standard color maps, this ensures that perceptual saliency aligns with biological meaning, with no salient colors (e.g. red) arbitrarily assigned. Technically, this involves computing the cell type similarity matrix, embedding this matrix into 3D via multidimensional scaling (MDS) to best preserve distances, then interpreting this 3D space as the LUV axes of the perceptually uniform LUV color space. We released a Python package which computes such colormaps called `ColorMyCell` to accompany this manuscript (***Benjamin, 2025***).

This dataset also contains labels of the brain area of each cell. These are inherited from the CCF coordinate space. We used these annotations as training data when predicting area from cellular transcription. As downloaded, these areas were at the finest level of the hierarchical tree of brain annotations and distinguished between cortical layers. In order to obtain the cortical area without regard to layers, we ascended one level up the hierarchical ‘ancestor’ tree of annotation layers in the Allen Brain Atlas. The resulting areas, displayed in e.g. ***Figure 3***, comprise 42 functionally distinct cortical areas.

#### Data description: COVID-19

We downloaded three datasets from CellXGene containing single-cell RNA sequencing data from healthy controls and patients of varying infection severity. The COMBAT Consortium dataset contained 637,266 cells from 85 donors sequenced with 10x 5’ v1 technology (10 healthy, 29 mild COVID-19, and 46 severe COVID-19) (***COMBAT, 2022***). The Ren et al. dataset contains 1,456,806 cells from 185 donors sequenced with 10x 3’ v3 and 10x 5’ v2 technologies (25 healthy, 77 mild COVID-19, and 83 severe COVID-19) (***Ren et al., 2021***). The Stephenson et al. dataset contains 585,153 cells from 106 donors sequenced with 10x 3’ transcription profiling (23 healthy, 55 mild COVID-19, and 28 severe COVID-19) (***Stephenson et al., 2021***). All three datasets were downloaded from CellXGene and contained pre-computed quality control filtering. Samples were labeled by patient condition at the time of blood draw.

To standardize labels across datasets, we mapped each dataset’s original severity annotations to a common schema of *control, mild*, and *severe*. In the COMBAT dataset, `COVID_SEV` and `COVID_CRIT` were mapped to *severe*, while `COVID_MILD` and `COVID_HCW_MILD` were mapped to *mild*. In the Ren et al. dataset, `severe/critical` and `mild/moderate` were mapped to *severe* and *mild*, respectively. In the Stephenson et al. dataset, `Severe` and `Critical` were grouped as *severe*, and `Mild` and `Moderate` as *mild*; samples labeled as `asymptomatic, non-COVID`, or `LPS-stimulated` were excluded. Donors with fewer than 1,000 cells were removed from all datasets. Because the severity classes are imbalanced across datasets, we trained all models with a frequency-balanced objective and report balanced accuracy (the accuracy computed within each class, then averaged across classes).

#### Tokenization

All cells’ mRNA transcript vectors were ‘tokenized’, or converted into a format readable by Transformer models, with the following procedure. We used an identical strategy as performed by Geneformer, but wrote custom code to accept data in anndata format (***Virshup et al., 2023***). First, each cell was normalized by the total counts in each cell. All mRNA counts were then normalized by a fixed vector representing the median per-gene counts in a reference data corpus. When fine-tuning Geneformer, it was empirically better to use the same median count vector used to train Geneformer (i.e. on human data) rather than re-establish median counts in mouse data. These normalized counts were then rank sorted, and the indices of the sort (i.e. the output of argsort) was used as the model input. For example, a model input of [gene 5, gene 2, …] describes that the gene 5 had highest relative transcription, followed by gene 2, and so on. In a modification of the Geneformer pipeline, we ensured genes with zero counts were given a special token `<pad>`. We also modified the Geneformer pipeline to prepend a `<cls>` token. The ‘sentences’ seen by the model were thus of the form [<cls>, gene 5, gene 2, …, <pad>]. The tokens are mapped to high-dimensional embedding vector via a dictionary learned within TissueFormer.

#### Group construction strategy

In order to train and predict with TissueFormer, it is necessary to prepare groups of cells which share a label. This requires code infrastructure to appropriately group cells according to the proper criteria. Since single-cell datasets are already quite large, we took an online strategy which loaded the single-cell dataset (after tokenization) and constructed groups online via a custom sampler and data collator within the Hugging Face code ecosystem for training TissueFormer.

For the COVID-19 datasets, groups of *N* cells are sampled uniformly at random from each donor. For brain annotation, our strategy was to select cells in groups which are approximately cortical columns, reflecting the columnar organization of the cerebral cortex in which neurons within a radial column share functional properties (***Mountcastle, 1997***). In order to cover the cortical surface with a roughly equal density, our pipeline first selected a seed cell at random from the training data, which potentially covers multiple brains. We next extracted the *N* cells in the same brain which were closest to that cell in the XY plane of the flatmap coordinate system. This resulted in a group of *N* cells which lie within a column. The width of the column differed in each group depending on the local density of cells. Note that because each brain contains a comparable number of cells, the column size in the test brain was comparable to the column size during training despite there being 3 training brains and 3 times the overall number of cells.

To test whether the increase in accuracy with group size was driven by cell type diversity rather than by averaging away measurement noise, we also constructed homogeneous groups restricted to a single cell type. For each group, a seed cell was selected at random and the *N* nearest cells of the same type were chosen, using the second level of the cell type hierarchy (“H2” types from ***Chen et al. (2024)***). This removes between-type diversity while preserving within-type variation. A separate TissueFormer model was trained from scratch on these homogeneous groups for each cell type.

### Predicting area from single cells

To provide a baseline for the multi-cell methods, we trained several machine learning models to predict area from single-cell data. Benchmark models either saw cell type (H3 types), raw transcriptomic data, or both in the case of the k-nearest neighbor cell type model. To finetune Murine Geneformer, we removed the last layer and replaced it with a linear classifier of area. We found best performance with a progressive training strategy popular in supervised finetuning. For one epoch, we froze all but the last layer, then each epoch progressively unfroze one Transformer layer in Murine Geneformer. We used a batch size of 32 and learning rate 0.0005. Results of training this model from scratch without pretraining are visible in ***Figure 2—figure Supplement 1***e.

### TissueFormer training

For the brain annotation task, we used leave-one-brain-out cross-validation: in each of four folds, three brains were used for training (90% of cells) and validation (10% of cells from the same three brains), and one brain was held out entirely as the test set. Random seeds were fixed across all methods to ensure identical data splits.

We trained TissueFormer on an NVIDIA H100 card with a batch size of 4096 total cells divided into *N* groups, each with |*G*| = 4096/*N* cells. A single model trains to completion for this dataset set (1.4 million cells) in 10 epochs in less than an hour. The most efficient training strategy involves saturating the GPU memory, and this is largely driven by the overall number of cells rather than the number of distinct groups. We trained all models with varying group size *N* to convergence to ensure a fair comparison. However, as convergence can be difficult to verify, we attempted to further match the training resources for each model by comparing models with an equal number of training steps, which is proportional to the number of cell groups seen.

Due to the imbalance in the size of brain areas, some classes have many more cells than others. A Bayes optimal model trained on this data would learn an unequal prior over classes, adding a slight bias to classify cell groups as belonging to the largest areas. This may not be desirable in certain applications of brain mapping. For example, cells on the boundary between a large area and a small area which have zero evidence towards one or the other will be classified into the large area due to the prior. This would cause an inflation of large areas. In ***Figure 3*** we display the results of a model trained with an objective designed to eliminate this bias and enforce an equal prior. Specifically we down-weighted examples from large areas with the factor as computed by the `compute_class_weight` function in scikit-learn (***Pedregosa et al., 2011***). For *M* total cells, *C* classes, and *M*_*i*_ cells in class *i*, this factor is 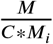. This ensures that the magnitude of the loss function equally reflects examples from all areas.

### Pseudobulk and type composition benchmarks

To ensure a fair comparison with TissueFormer, we constructed train and test sets using the same data loading pipeline with the same random seed. For the pseudobulk results, we then averaged the mRNA transcript counts within each group of cells, then trained our benchmark methods. For the cell type composition models, we extracted the cell type of each cell in the group at the finest available level of the cell type hierarchy, then constructed a normalized histogram reflecting the relative composition of cell types in that group. Each bin of this histogram corresponds to one cell type, and its value is the fraction of cells in the group belonging to that type, such that the histogram sums to one. For both data modalities, our benchmark methods were logistic regression and random forests. Specifically, we trained logistic regression with the lbfgs solver. We used nested cross-validation to determine the regularization parameter, and found that no regularization was optimal at this dataset size. We also used a random forest classifier with 200 estimators, max depth of 15, 33% of features seen per tree, and a minimum samples per split and 5 and per leaf at 2. We used the scikit-learn implementation of both classifiers (***Pedregosa et al., 2011***).

### Brain map visualization

To create ***Figure 3***a, we first trained 4 models in a cross-validated scheme, with 1 brain held out each fold. To achieve a high-resolution coverage, we then predicted on the test brain using every cell in the test brain once as a center seed of group selection. The prediction on each cell was stored as its label. These are visualized in ***Figure 3—figure Supplement 1***. We then constructed a method to present smooth brain maps with clear boundaries between areas. Namely, we trained a support vector classifier (SVC) with a radial basis function kernel on the cells and their predicted labels, then visualized the predictions on pixels. We used the cuML implementation, a GPU-accelerated method, and used *γ* = 0.00001 and *C* = 1 as hyperparameters of the SVC (***Raschka et al., 2020***). Intuitively, this method acts to color each pixel in ***Figure 3***a by a weighted average of the TissueFormer-predicted labels of the cells surrounding that pixel. The weights are selected due to a combination of proximity to the pixel and a factor determined by the algorithm to minimize the rate at which cells are misclassified, i.e. located on a pixel with a different color (area) than that cell’s TissueFormer-predicted area. To mask regions with low cellular density, we estimated the spatial density of cells on the flatmap using a Gaussian kernel density estimator (bandwidth 12, cuML implementation). The resulting density was normalized to [0, 1], and pixels with normalized density below 0.04 were overlaid with a semi-opaque white mask (95% opacity). To convert predicted area sizes from flatmap pixels to physical units (mm^2^), we estimated the physical cortical surface area represented by each flatmap pixel. Because the butterfly flatmap is a distorted 2D projection of the curved cortical surface (***Wang et al., 2020***), pixels in different parts of the flatmap correspond to different amounts of cortical surface. We computed this correction by mapping each flatmap pixel back to its corresponding coordinate on the pial surface (at 10 *μ*m resolution) and estimating the local surface area element via finite differences between neighboring pixels. The area of a predicted region was then the sum of these per-pixel surface areas over all pixels classified as that region.

### COVID-19 severity prediction training

For the COVID-19 severity prediction task, we used 5-fold stratified group *k*-fold cross-validation, where all cells from a single donor were assigned to the same fold and folds were stratified by severity label. Within the training folds of each split, 10% of donors were held out for validation, again stratified by severity. We trained TissueFormer with a learning rate of 5 × 10^−5^, a batch size of 512 total cells divided into groups of size *N*, and the AdamW optimizer. Training used early stopping with a patience of 3 epochs, monitoring balanced accuracy on the validation donors. We used mixed-precision (fp16) training for group sizes greater than 1. As with the brain task, we trained models with a frequency-balanced objective and report balanced accuracy to account for class imbalance across severity levels.

### Deep learning benchmarks

To compare TissueFormer against other end-to-end deep learning methods for multi-cell classification, we benchmarked three recent architectures: CellCnn (***Arvaniti and Claassen, 2017***), scAGG (***Verlaan et al., 2025***), and ScRAT (***Mao et al., 2024***). For each method, we matched the number of cells per group to the group size *N* used for TissueFormer and trained and evaluated on identical data splits. Hyperparameters for each method were taken from the original publications.

CellCnn applies a 1D convolution (kernel size 1, 6 filters) over cells followed by top-*k* mean pooling (top 5% of cells per filter) and a linear classifier. We trained with Adam (learning rate 0.01 for brain, 0.001 for COVID), batch size 200, dropout 0.5, L1 and L2 regularization of 10^−4^, and early stopping (patience 5 for brain, 15 for COVID). Inputs were z-score normalized.

scAGG uses a two-layer MLP (input → 1024 → 512, ELU activations) followed by mask-aware mean pooling and a linear classifier. We trained with Adam (learning rate 0.001), batch size 8, dropout 0.1, weight decay 5 × 10^−4^, and 2 epochs. Inputs were normalized to counts per 10,000 and log-transformed. For the COVID datasets, we selected the top 1,000 highly variable genes.

ScRAT employs a Transformer encoder (1 layer, 8 attention heads, hidden dimension 128, feedforward dimension 256) over cell embeddings produced by a two-layer MLP (input → 256 → 128). Cell representations are mean-pooled and passed to a classification head. We trained with Adam and a cosine learning rate schedule with 10-epoch warmup (peak learning rate 0.003), batch size 16, dropout 0.3, weight decay 10^−4^, and early stopping after epoch 50 with patience 2. All methods were trained with class-balanced objectives.

### Leave-one-cell-type-out analysis

To investigate which cell types drive TissueFormer’s predictions, we performed leave-one-cell-type-out (LOO) perturbation experiments. For each cell type present in a group, we removed all cells of that type and re-evaluated the model, measuring the change in cross-entropy loss relative to the unperturbed group. A large increase in loss upon removal indicates that the cell type is important for the correct prediction. We report results broken out by predicted label, averaging across all groups sharing that label. For the COVID-19 task, LOO analyses were performed using the *N* = 32 model on the COMBAT dataset.

## Results

We evaluated TissueFormer on two tasks in which cellular diversity is expected to be informative: predicting cortical area identity from spatial transcriptomic data in mouse brains, and predicting COVID-19 severity from single-cell RNA sequencing of peripheral blood. For the brain annotation task, we compared TissueFormer against traditional metrics such as cell-type ratios and pseudobulk expression, as well as other deep learning methods. TissueFormer outperformed single-cell models on both tasks, confirming the practical value of attending across cells when predicting population-level phenotypes. In the brain task, we additionally found evidence that standard tissue alignment procedures do not correctly register tissue into the correct functional areas.

### Single-cell profiles are not informative of cortical area

The utility of attending across cells can be made clear by comparing with the performance of machine learning methods which are trained to predict the area given only a single cell. To investigate this, we first split the data into training data (90% of cells in 3 brains), validation data (the remaining 10% of the same 3 brains), and a held-out test brain (***Figure 2***b). We then trained a range of methods on the task of predicting the cortical area of a single cell from its transcriptome.

**Figure 2.**
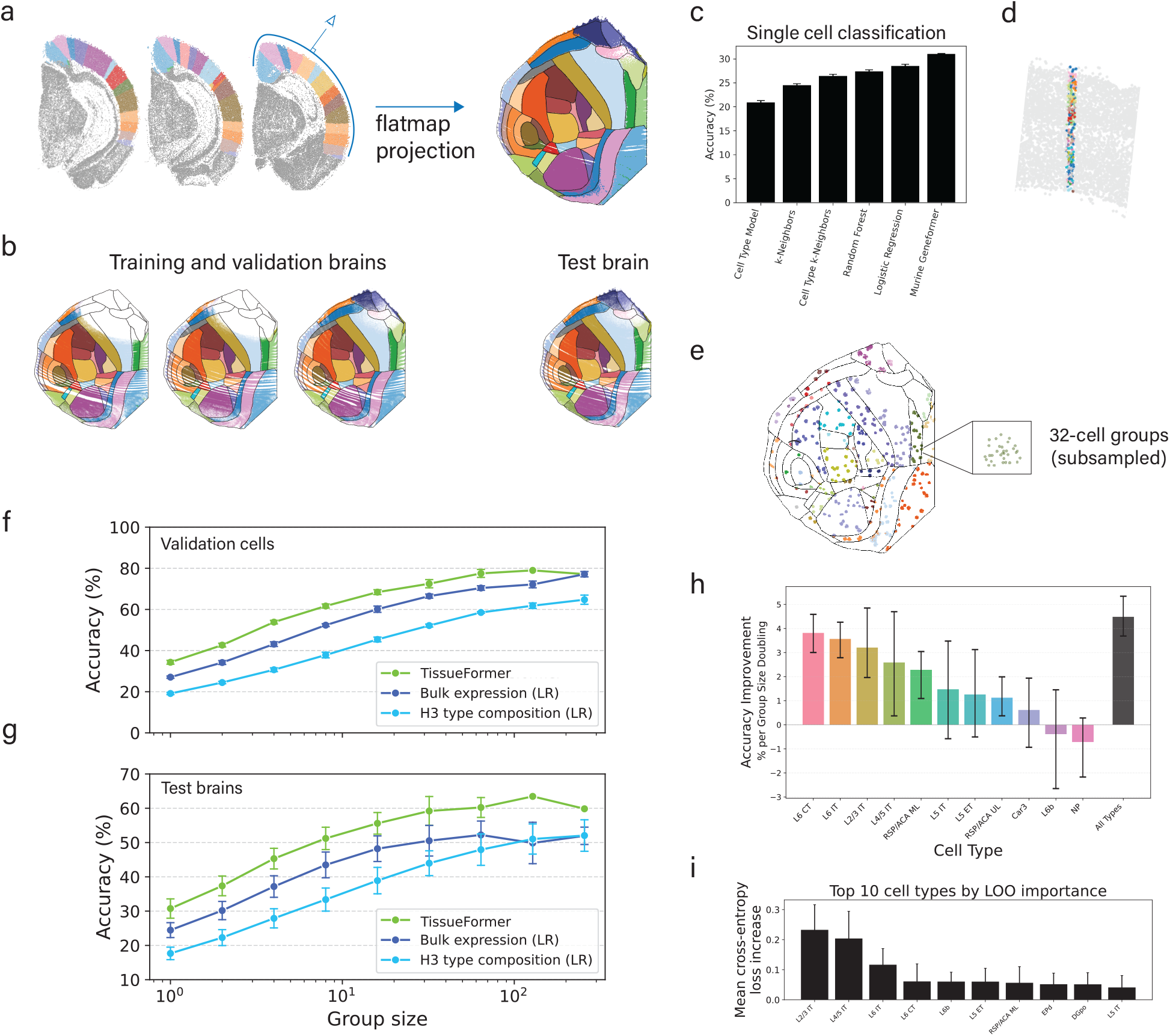
Comparison of accuracy at predicting brain regions shows that TissueFormer outperforms other methods. **a)** Cortical cells from coronal slices of one hemisphere (left, colored by area) are visualized as a ‘flatmap’, a top-down projection. Color legend is in ***Figure 3*.b)** Models were evaluated using 4-fold crossvalidation with 3 or 4 brains used for training and validation labels, 1 of 4 brains held out for testing. Shown here are true labels for each cell. **c)** Comparison of accuracy when predicting the area of a single cell in a test brain from its transcriptome shows that all methods perform poorly, though with Murine Geneformer performing best. **d)** An example cell group (*N* = 256) plotted in slice coordinates, colored by cell type. **e)** A sample of groups (*N* = 32) plotted in flatmap coordinates, here colored by the modal area of the group. **f)** Validation accuracy as a function of group size shows that TissueFormer outperforms logistic regression (LR) given either pseudobulk transcription or a histogram of cell types. **g)** Test brain accuracy. Error bars in f-g represent standard deviation across 4 folds. **h)** The average rate of test accuracy improvement with group size for the above TissueFormer (black bar) compared to the accuracy curve slopes of several TissueFormer models trained on homogeneous groups containing only a single cell type. Accuracy curves are visible in supplement 1. **i)** Average leave-one-cell-type-out (LOO) importance across all predicted areas. Each bar shows the mean increase in loss when all cells of that type are removed from a group. Cell types are ranked by overall importance. **Figure 2—figure supplement 1**. Effect of pretraining on TissueFormer, performance of random forest and deep learning benchmarks, and controls for unequal area size and sampling density. **Figure 2—figure supplement 2**. Leave-one-cell-type-out analysis for brain annotation.

The methods tested ranged in complexity from simple heuristics to modern single-cell foundation models (***Figure 2***c). We included three standard machine learning benchmarks, namely, a logistic regression model, a random forest model, and k-neighbors classifier. These models map the vector of mRNA transcript counts to the label after a log(1 + *x*) transformation and z-scoring, and their hyperparameters were tuned on the validation set. We also trained two models which make use of the cell types of the cell in question. These cell types were previously constructed at three levels of granularity (***Chen et al. (2024)***); we selected the finest level. The ‘cell type model’ predicts the area of a cell based on the most common area of cells of the same type in the training set. The ‘cell type k-neighbors’ model labels the target cell based on the area of the k-closest neighbors with the same cell type in the training set. Finally, we fine-tuned Murine Geneformer to predict the area of each cell.

Despite the diversity of modeling approaches, all six models performed poorly, with classification accuracies ranging from 20–31%. This highlights the challenge of predicting cortical area from single-cell transcriptomes, consistent with previous findings that many cell types are broadly distributed across the cortex (***Tasic et al., 2018; Yao et al., 2021b; Chen et al., 2024***). While some types are more localized and thus were predicted with higher accuracy (for example, RSP/ACA or Retrosplenial/Anterior Cingulate excitatory cells), overall performance was low. Among the models tested, the Transformer-based Murine Geneformer achieved the highest accuracy, modestly outperforming the others. This is consistent with recent results showing strong performance of large pretrained models on downstream tasks (***Theodoris et al., 2023***). These results motivated our turn toward models that integrate information across cells.

### Annotating cell groups with TissueFormer

To apply TissueFormer, we first created a pipeline to group cell sharing a label. Specifically, we grouped cells into cylindrical columns which fall inwards from the surface of the cortex (***Figure 2***d-e). Note that this strategy can easily be generalized to other shapes for spatial data from other tissues, or to other sampling strategies for non-spatial data. The cerebral cortex can be viewed as a layered cake draped over the exterior of the mouse brain, with functional areas as slices (***Figure 2***a). Thus, cortical columns contain cells in the same functional area. Our pipeline programmatically selects a single cell, then grows a cylinder centered on that cell up to the minimum radius needed to contain *N* total cells. The density of this dataset is such that a group of 64 cells spans on average 0.008 mm^2^, equivalent to a 56 µm radius. Each columnar group of *N* cells is labeled with the most common CCF annotation of cells in that group.

We next trained TissueFormer to predict the group’s label from the set of transcriptomic profiles within the group. We again cross-validated by training on 90% of cells from three brains, validating on the remaining 10% of cells, and testing on one held-out brain. As we varied the number of cells in each group from *N* = 1 to *N* = 256, we observed that the validation accuracy of TissueFormer increased from around 35% to nearly 80% accuracy (***Figure 2***f). On test brains, the accuracy Tissueformer also increased markedly with group size from 30% to over 60% accuracy (***Figure 2***g). Providing the cortical depth of each cell as additional input to the model did not improve results, suggesting that depth represents redundant information (***Figure 2—figure Supplement 1***f). Note that random guessing on this 1-of-42 classification task yields 8% accuracy due to imbalanced classes. Similar relative accuracies were seen when training with a class-balancing weighted objective (***Figure 2—figure Supplement 1***c). These results demonstrate that TissueFormer is able to dramatically outperform single-cell models at predicting tissue identity.

As the group size increased from *N* = 1 to *N* = 256, the accuracy increased smoothly and roughly logarithmically with group size, indicating a predictable scaling law with more cells. Performance saturated at *N* = 128 cells, possibly because at this scale and at this density of cells per brain the groups sometimes straddle boundaries, at which point group identity begins to lose meaning. Indeed, at *N* = 256 over 70% of groups contained a boundary (***Figure 2—figure Supplement 1***g). Higher sampling densities would enable larger group sizes in smaller locales, and thus possibly an even higher saturating accuracy. Due to this problem of spatial density, it is not possible to verify the maximum group size at which cells contain no further information about area. Overall, the considerable increase of performance with group size highlights the importance of the ability to integrate and compare information across single cells.

The cell embedding module in the above experiment was previously pretrained on a masked mRNA transcript prediction task on several other datasets containing tens of millions of human and mouse cells. In principle, this pretraining step could help, hinder, or have little effect on our current supervised task of predicting brain area. To investigate its impact, we also trained a TissueFormer model from a random initialization without any pretraining. Surprisingly, the performance of this randomly initialized model was indistinguishable from the pretrained model for all group sizes (***Figure 2—figure Supplement 1***d). This is likely due to the large number of cells in our labeled dataset. To confirm this, we varied the number of training cells used to train both a pretrained and a randomly initialized TissueFormer from one thousand cells to the full dataset of over two million. We found that pretraining indeed offered an advantage, but only in the intermediate range of less than one million cells (***Figure 2—figure Supplement 1***e). Thus, this spatial transcriptomic dataset is large enough that transfer learning from Murine Geneformer offers no advantage.

We next compared the performance of TissueFormer with two standard methods for processing transcriptomic data of groups of single cells. We first examined pseudobulk analyses, which average the single-cell profiles in the group to approximate the measurements of a traditional bulk sequencing experiment. Logistic regression trained on pseudobulk vectors underperformed TissueFormer, yet still showed a nearly logarithmic increase of accuracy with group size (***Figure 2***f). A random forest model trained on the same data underperformed logistic regression (***Figure 2—figure Supplement 1***b). Next, we trained logistic regression and random forests to predict area from the cell type composition of a group, which was represented as a histogram of cell types. We used the finest-grain categorization of cell types from ***Chen et al. (2024)*** for this analysis. Overall, type composition was less informative of area than pseudobulk expression (***Figure 2***f). Intriguingly, each method displayed a logarithmic increase in accuracy with group size despite the differences in their representations of RNA transcription. We also compared TissueFormer against three end-to-end deep learning methods for multi-cell classification: CellCnn (***Arvaniti and Claassen, 2017***), scAGG (***Verlaan et al., 2025***), and ScRAT (***Mao et al., 2024***). TissueFormer outperformed all three across all group sizes (***Figure 2—figure Supplement 1***i).

The logarithmic increase in accuracy with group size across all methodologies could in principle be driven by two effects. One possibility is that the diversity and particular composition of cells provided a new signal not available to single cell models. An alternative possibility is that cells contained independent measurement noise which can be averaged away. Some evidence to the first possibility is that benchmarks based on cell type composition alone showed such an increase (***Figure 2***f-g). To further distinguish these factors within TissueFormer, we reasoned that we could artificially restrict the diversity of cells in each group and then re-test a method’s performance. If the scaling were due to measurement noise alone, then this would have little effect and performance would likewise increase with a similar slope on a log-linear plot. We therefore constructed a new method of constructing groups such that they contain only a single cell type from the second level of granularity in the cell type hierarchy (“H2” types). Note that there is still significant diversity within each H2 type, albeit much less than the general population. After training a TissueFormer model from scratch on these homogeneous groups, we found that the test set accuracy given a group of homogeneous type scaled more slowly with increasing group size *N*, on average (***Figure 2***h). Groups of cells of certain types such as NP cells showed little improvement with larger size. Other types, such as L6 IT cells, nevertheless show comparable increases, likely indicating higher and more informative diversity in that population. The areas of some cell types are more easily predictable than other types due regional localization (***Figure 2—figure Supplement 1***h). These analyses confirmed that the increase in accuracy with group size was at least partially driven by comparative signals across a cell group rather than solely by averaging away technical noise.

To understand which cell types drive TissueFormer’s predictions, we performed leave-one-cell-type-out (LOO) experiments in which all cells of a given type were removed from each group and the change in loss was measured (***Figure 2***i). When broken out by predicted area (***Figure 2—figure Supplement 2***), removing L4/5 IT cells caused the largest loss increase in somatosensory areas, while L2/3 IT cells were most important for visual areas. These patterns are consistent with known regional variation in laminar cell type composition across the cortex.

The decrease in accuracy between validation and test brain accuracy could have arisen due to multiple reasons. A first reason is the differences in spatial coverage of data from each brain, visible in ***Figure 2***b, which may cause test brains to contain areas not in the training data. However, correcting for this effect by only testing data with high density in the training set yielded improvements in accuracy of at most a few percentage points (***Figure 2—figure Supplement 1***a). Secondly, this could have reflected other batch effects across brains affecting single-cell measurements, effectively making this test brain an out-of-distribution test for the methods. Finally, it is also possible that ‘errors’ in classification were due to actual changes in neuroanatomy which were correctly predicted by the model but were mislabeled by the pipeline of registering each brain to the CCF. We investigate this possibility in the following section.

### Predicted cortical maps

TissueFormer can be used as an automated pipeline to create maps of cortical anatomy from single-cell transcription after being trained on reference brains. Note that annotating from reference brains is a supervised task, not to be confused with the unsupervised task of spatially clustering single-cell data (see Discussion). In ***Figure 3***a, we demonstrate this capability and display the predicted cortical maps for each of 4 mice treated as a held-out test set. Pixels in these maps were colored by a weighted average of nearby cells’ predictions (see Methods). For visualization we used the predictions of TissueFormer trained with a class-size-balancing objective ensuring that the prior probability was uniform across all areas regardless of area size. We found that all four maps were coarsely similar to the reference annotation (see ***Figure 2***a), supporting the finding in ***Figure 2*** that TissueFormer is able to predict brain area more accurately than other approaches.

**Figure 3.**
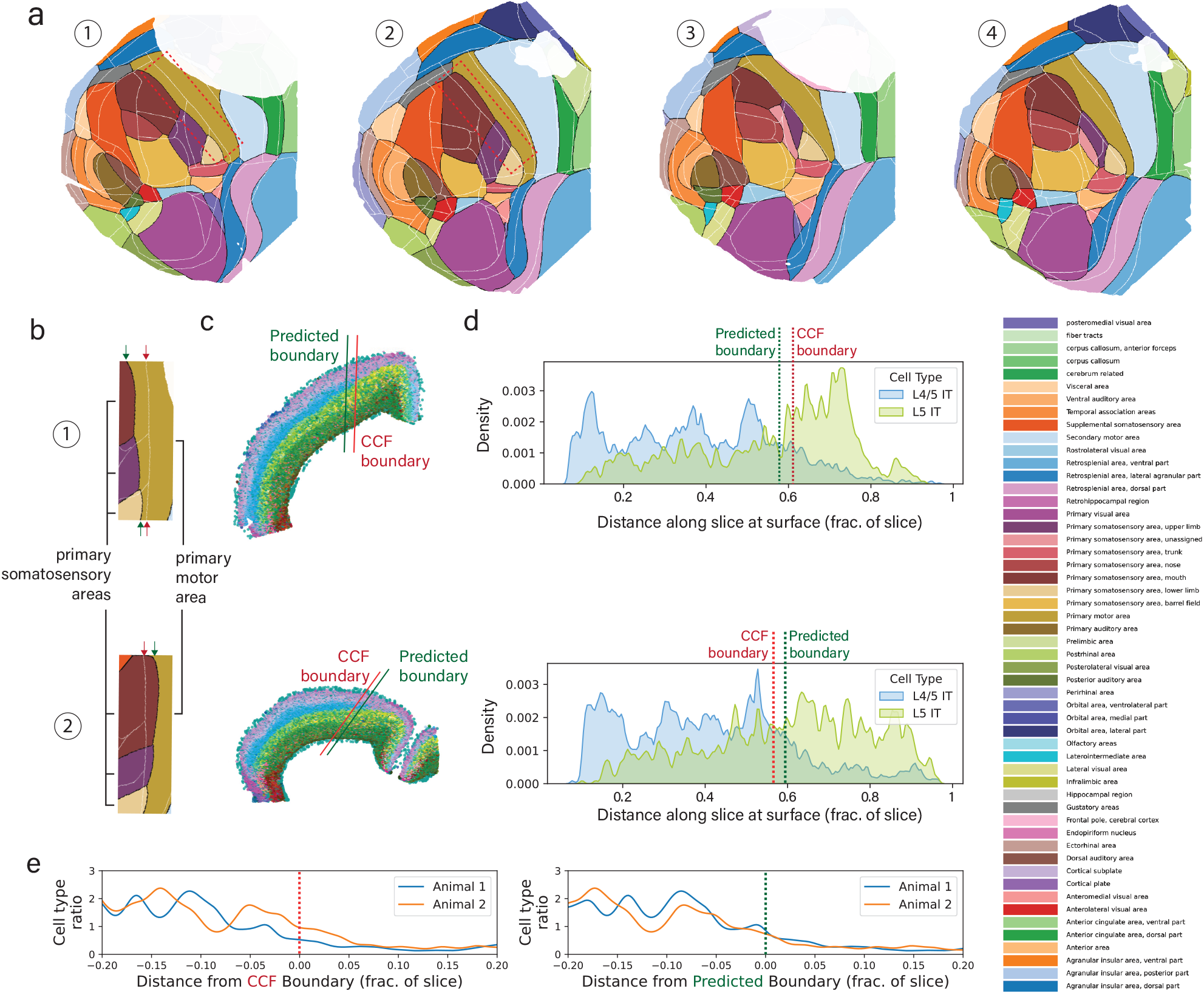
Creation of cortical maps from spatial transcriptomics using TissueFormer. **a)** Predicted cortical areas for each brain when held-out as a test brain. Predicted boundaries are in black, and the reference Common Coordinate Framework (CCF) boundaries are overlaid in white. Areas with low cellular density are masked in white (see Methods). **b)** In brains 1 (top) and 2 (bottom), the somatosensory-motor boundary is displaced relative to the CCF, but in opposite directions. **c)** Example slices from brains 1 and 2 which contain the somatosensory-motor boundary boundary. Points are cells, colored by cell type with similar cell types assigned similar colors (see Methods). **d)** Spatial density along the slices of two cell types, L4/5 IT and L5 IT, in the same colors and slices as (c). **e)** The cell type ratio (L4/5 IT / L5 IT) aligned to the CCF boundary (left) or predicted boundary (right), showing an inter-animal shift in CCF vs. inter-animal consistency in the predicted maps. **Figure 3—figure supplement 1**. Predicted areas of single cells, highlighting discrepancies (‘errors’) with CCF labels. **Figure 3—figure supplement 2**. Comparison to maps of mRNA transcription of single genes.

A close inspection of these cortical maps revealed several notable differences in the predicted anatomy of each mouse. For example, the boundary between the somatosensory cortex and the motor cortex is consistently shifted in the posterior-lateral direction in brain 1 relative to the Common Coordinate Framework (CCF) annotations, yet is consistently shifted in the opposite (anterior-medial) direction in brain 2 (***Figure 3***b). Additionally, the primary visual area is shifted medially in brain 2, yet laterally in brains 3 and 4. Likewise, the primary auditory area is shifted medially in brain 2 yet is expanded and shifted laterally in brain 4. These shifts are a predominant cause of why test brain ‘accuracy’ in ***Figure 2***g is lower than validation accuracy.

While it is possible that the discrepancies between the CCF annotation and the predicted annotation are ‘errors’ of the model, it is also possible that this reflects true differences in anatomy between mice or errors in registration with the CCF. The CCF represents the average anatomy of over 1,000 mouse brains, obscuring potential individual variability between mice. To investigate this possibility, we examined the somatosensory-motor boundary in more detail. This boundary can be canonically identified by differences in cell type composition. In particular, motor cortex is classically considered to lack a Layer 4 (***Brodmann, 1909***), and thus should show a large decrease in the density of Layer 4/5 IT excitatory cells relative to Layer 5 IT cells. This easily verifiable boundary was qualitatively visible in coronal slices colored by cell type (***Figure 3***c). To confirm its location, we examined the density of these two cell types along slices taken from brain 1 and brain 2 which intersect the sensory/motor boundary (***Figure 3***d). We found that the CCF boundary was inconsistent across brains relative to the ratio of cell types, but TissueFormer’s predicted maps showed a consistent 1:1. Furthermore, in the CCF boundaries, we observed a large shift between mice in the ratio for large distances on either side (***Figure 3***e). In contrast, the density ratios were consistent in their alignment to the predicted boundaries. Thus, a closer inspection of cell type distributions around this verifiable boundary revealed a closer alignment with our predicted atlas than with the boundaries due to CCF alignment.

To provide further verification of the predicted cortical maps, we examined the spatial distributions of individual genes known to correlate with area identity (***Figure 3—figure Supplement 2***). For example, Tshz2 forms a striking medial-to-lateral gradient with high abundance in anterior- and posterior-cingulate/retrosplenial areas with steep drops at the boundaries to motor, somatosensory and visual fields (***Yao et al., 2021b***). Across all four brains, the inter-animal differences in Tshz2 aligned better with the predicted boundary than the CCF area boundaries. Other genes which are also selectively expressed in retrosplenial areas, such as as Coro6 and Zfpm2 (***Chen et al., 2022***), shared this pattern, as did genes with selectively expressed in the dorsal but not the ventral retrosplenial area, such as Nell1 and Zmat4 (***Hashikawa et al., 2020***). Meanwhile, the sensory/motor border analyzed in ***Figure 3*** was correlated to the expression density of Rorb1, Brinp3, and Rcan2 (***Cederquist et al., 2013***), all of which showed consistent shifts across animals with the predicted maps but not with CCF boundaries. These examples of genes with clear spatial boundaries in their patterns of expression lend support to the predicted maps’ veracity.

The BARseq dataset also includes four brains from mice that were binocularly enucleated (both eyes surgically removed) at birth, each paired with a control littermate. Removing visual input during development is known to shift the cell-type composition of visual areas toward that of neighboring cortical areas (***Chen et al., 2024***), raising the question of how this reorganization manifests at the level of predicted area maps. We applied TissueFormer, trained on the four control brains, to the four enucleated brains and generated predicted cortical maps for each (***Figure 4***). The maps were consistent across the four enucleated brains. The predicted area of primary visual cortex (VISp) was significantly smaller in enucleated brains (mean 7.1 mm^2^) than in controls (mean 9.0 mm^2^; paired *t*-test across littermates, *p* = 0.01, *n* = 4 pairs). These automated maps illustrate how TissueFormer can be used to quantify the effects of developmental perturbations on cortical organization across individual brains.

**Figure 4.**
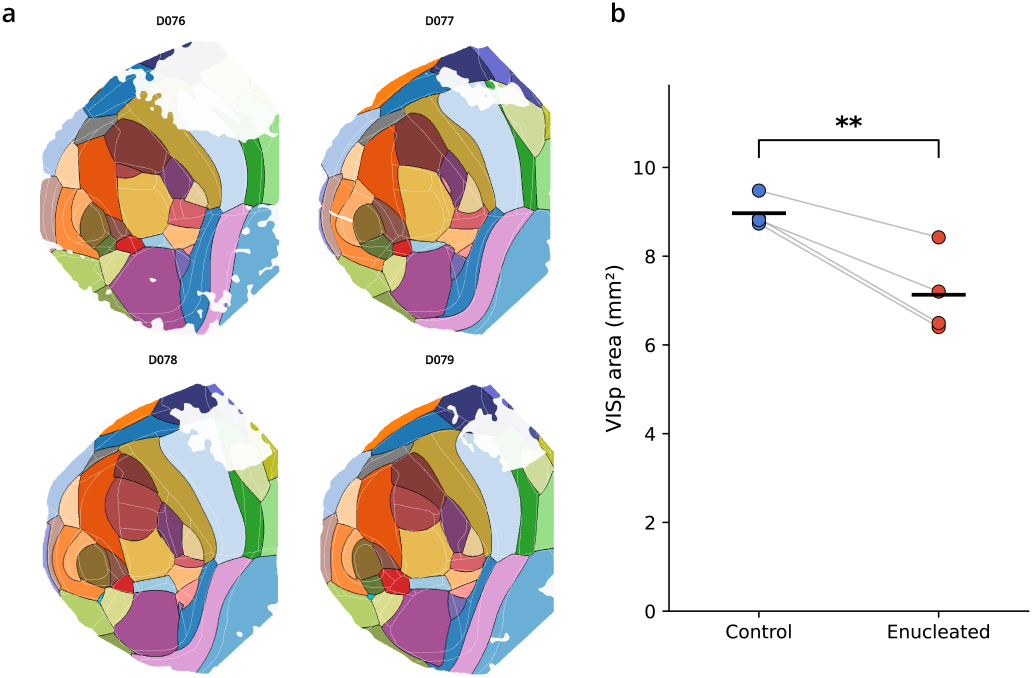
TissueFormer detects reduced visual cortex area in mice deprived of visual input during development. Predicted cortical maps were generated for four mice binocularly enucleated at birth, each paired with a control littermate, using TissueFormer trained exclusively on the four control brains. Maps are constructed identically to ***Figure 3***a: predicted boundaries are in black, CCF boundaries are overlaid as thin white lines, and areas with low cellular density are masked in white (see Methods). The predicted area of primary visual cortex (VISp) was significantly smaller in enucleated brains (mean 7.1 mm^2^) than in controls (mean 9.0 mm^2^; paired *t*-test across littermates, *p* = 0.01, *n* = 4 pairs).

### Predicting COVID-19 severity from blood transcriptomics

To test whether TissueFormer generalizes beyond spatial brain data, we applied it to a fundamentally different prediction task: classifying COVID-19 infection severity (control, mild, or severe) from single-cell RNA sequencing of peripheral blood. This task differs from brain annotation in several important ways. The data are non-spatial, so cells are grouped by donor rather than by spatial proximity. The organism is human rather than mouse. The sequencing modality is droplet-based 10x rather than probe-based BARseq. Finally, the phenotype is a clinical disease state rather than an anatomical label. We compiled three independent cohorts totaling over 2.6 million cells from 376 donors: COMBAT (***COMBAT, 2022***), Ren et al. (***Ren et al., 2021***), and Stephenson et al. (***Stephenson et al., 2021***). Each dataset was evaluated independently using 5-fold stratified group *k*-fold cross-validation with all cells from a donor assigned to the same fold (see Methods).

As with brain annotation, we varied the number of sampled cells per group and compared TissueFormer against pseudobulk and cell type composition baselines (***Figure 5***a, left column). Across all three cohorts, TissueFormer’s balanced accuracy increased with group size and exceeded that of logistic regression and random forest models trained on either pseudobulk expression or cell type histograms. The advantage was most pronounced on the largest dataset, Ren et al., where at *N* = 64 TissueFormer achieved 70% balanced accuracy compared to 54% for the best classical baseline (logistic regression on cell type histograms) and 52% for random forest on pseudobulk. This gap persisted at *N* = 256 (69% vs. 58%). A similar pattern held when all three cohorts were combined. On the smaller COMBAT dataset, TissueFormer and logistic regression on pseudobulk were closer in performance (87% vs. 85% at *N* = 256), suggesting that the advantage of end-to-end learning is greatest when more donors are available for training.

**Figure 5.**
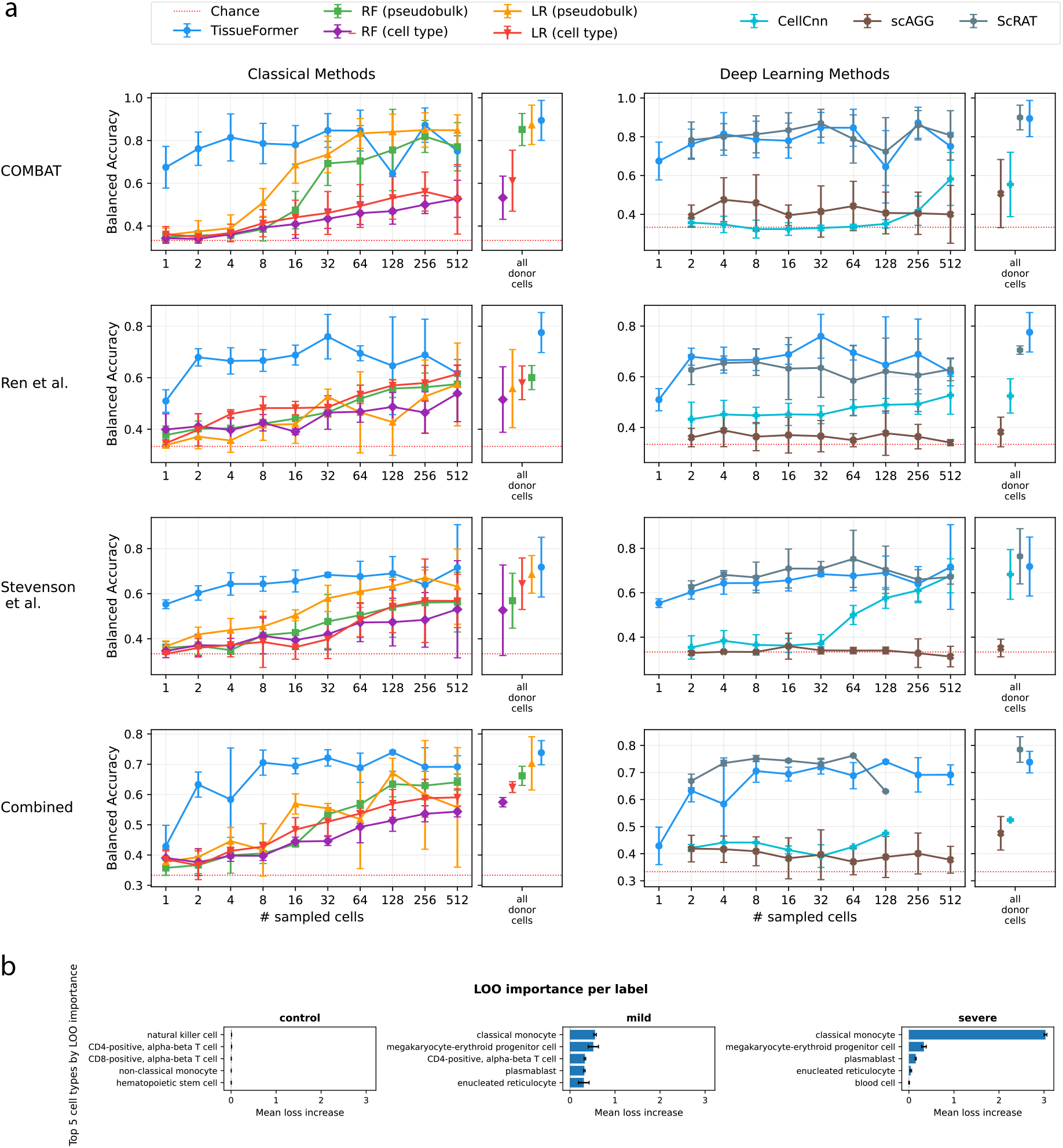
TissueFormer predicts COVID-19 severity from single-cell blood transcriptomics. **a)** Balanced accuracy as a function of group size for three independent cohorts (COMBAT, Ren et al., Stephenson et al.) and their combination. Left column: TissueFormer compared to logistic regression (LR) and random forest (RF) trained on pseudobulk expression or cell type composition histograms. Right column: TissueFormer compared to three end-to-end deep learning methods (CellCnn, scAGG, ScRAT). Error bars represent standard deviation across 5 folds. The condition “all donor cells” uses every cell from a donor as a single group for classical methods; for deep learning methods, which require a fixed group size, predictions are instead averaged over multiple groups of size *N* sampled from the donor. **b)** Leave-one-cell-type-out (LOO) analysis. Each bar shows the change in loss when all cells of a given type are removed from groups of a given severity class. Positive values indicate that the cell type contributes to correct classification; negative values indicate that its presence is distracting. Classical monocytes are the most important type for classifying severe samples.

We next compared TissueFormer against three end-to-end deep learning methods for multi-cell classification: CellCnn, scAGG, and ScRAT (***Figure 5***a, right column). The two Transformer-based methods, TissueFormer and ScRAT, consistently outperformed CellCnn and scAGG. scAGG remained near chance across all cohorts (33–40% balanced accuracy), and CellCnn was variable, performing near chance on COMBAT (42%) but reaching 61% on Stephenson et al. TissueFormer and ScRAT were not significantly different from each other: on individual cohorts the two methods achieved similar balanced accuracy (e.g. 87% vs. 86% on COMBAT, 69% vs. 61% on Ren et al. at *N* = 256), with ScRAT slightly ahead on the combined dataset (75% vs. 69%). These results indicate that Transformer-based architectures are well suited for multi-cell classification, substantially outperforming alternative deep learning approaches on this task.

To understand which cell types drive TissueFormer’s severity predictions, we performed leave-one-cell-type-out (LOO) experiments in which all cells of a given type were removed from each group and the change in loss was measured (***Figure 5***b). Classical monocytes were by far the most important cell type for correctly classifying severe samples, with their removal increasing the error rate on severe groups by 49%. For groups labeled mild, classical monocytes, megakaryocyte-erythroid progenitors, and CD4+ T cells all contributed to accurate prediction. No single cell type was critical for correctly identifying control samples. Interestingly, removing lymphoid types such as CD4+ or CD8+ T cells from severe groups actually reduced the loss, suggesting that these cells can be distracting when classifying disease severity. These patterns are consistent with the known expansion of monocytes and myeloid progenitors in severe COVID-19 (***COMBAT, 2022; Ren et al., 2021***).

## Discussion

We developed TissueFormer, a neural network architecture that attends to groups of single-cell transcriptomic profiles in order to predict a label or annotation common to that group. On a brain area classification task using multi-brain spatial transcriptomics, TissueFormer achieved 60% accuracy with groups of 64 cells on held-out test brains, compared to 30% for a matched single-cell model, and outperformed pseudobulk and cell type composition baselines. On a COVID-19 severity prediction task using non-spatial blood scRNA-seq from three independent cohorts, TissueFormer likewise outperformed classical baselines and matched or exceeded other deep learning approaches. In both settings, accuracy increased with group size, highlighting the importance of attending across cells when predicting tissue-level properties.

Because TissueFormer accepts arbitrary cell groupings, it can flexibly model phenotypes defined by anatomical contiguity (as in spatial transcriptomics), by patient sample (e.g., blood draws, biopsies), or by dynamic windows in longitudinal sampling. These applications include a wide range of use cases, from immunomonitoring in inflammatory disease to potentially the detection of cancer states (***Weinkauff et al., 1999; Sun et al., 2025***). As increasingly comprehensive healthy and disease cell atlases become available, population-aware models like TissueFormer offer a principled route to integrate those resources into predictive diagnostics that bridge cellular resolution and clinical decision-making (***Dann et al., 2023***).

TissueFormer provides a promising tool for automated brain mapping from spatial transcriptomics. Compared to the labels due to Common Coordinate Framework (CCF) registration, TissueFormer’s predicted labels displayed individual variability that better correlated with inter-animal differences in local gene expression and cell type distributions, despite being trained to predict CCF labels. This tool could thus expedite the study of individual differences in the size and location of transcriptomically-defined regions in the brain. As a concrete example, TissueFormer’s predicted maps revealed a significant reduction in the area of primary visual cortex in mice deprived of visual input during development, consistent with past analyses of cell-type compositional shifts in these brains (***Chen et al., 2024***).

Our approach to brain mapping is a supervised approach in which reference brains supply ground truth labels. Note that this is distinct from the approach of aligning tissue without regard to area labels, as in Tangram (***Biancalani et al., 2021***) and CAST (***Tang et al., 2024***). An important caveat of the supervised approach is the lack of perfect training data. No datasets of spatial transcriptomics are yet available which have animal-by-animal annotations of neuroanatomy verified by additional modalities such as projection tracing or functional imaging. Without access to such datasets, we instead trained to predict the imperfect CCF labels. In general, there is no formal guarantee that the predictions on held-out brains should reflect a better assessment than the CCF itself. While we are led by our empirical analysis to trust the model, this must be done with caution until a model can be trained on a dataset with multiple brains individually annotated with complementing experimental modalities.

In the current study, we did not take up the question of whether different boundaries than the Allen Brain Atlas would be in any sense better, as in recent spatial clustering methods. Current boundaries are widely used and important for shared nomenclature. However, the automated clustering and identification of spatially homogeneous regions within spatial transcriptomic dataset is an interesting and open research question. Several algorithms have already been developed for this purpose (***Dries et al., 2021; Zhao et al., 2021; Chitra et al., 2025; Hu et al., 2021; Dong and Zhang, 2022; Singhal et al., 2024; ?; Jackson et al., 2024***). Because these unsupervised methods discover spatial domains without reference labels, their outputs are not directly comparable to those of supervised methods like TissueFormer, which predicts known annotations from labeled training data. When algorithms from this family are applied to the mouse cortex, the resulting regions correspond more to cortical layers than to functional regions (***Ortiz et al., 2020; Partel et al., 2020; Lee et al., 2025***). This is consistent with there being larger differences in gene expression across layers than across cortical areas. In contrast, the functional specialization of neurons in cortex varies more across area than across layers within an area; neurons within a cortical column typically mediate the same cognitive tasks and furthermore respond to similar features within that task (***Mountcastle, 1997; Callaway, 1998***). Amid this dichotomy between transcriptomic and functional contiguity in space, we chose in this manuscript to classify functional areas for practical utility and for the reason of it being a benchmark task that leverages transcriptomic diversity within cellular ensembles. Nevertheless, it would be interesting in future work to train TissueFormer with self-supervised or contrastive objectives so as to discover brain areas *de novo*.

## Declarations

### Ethics approval and consent to participate

Not applicable.

### Consent for publication

Not applicable.

### Availability of data and materials

All code is available at https://github.com/ZadorLaboratory/TissueFormer. Brain spatial transcriptomic data are hosted on Mendeley Data (https://doi.org/10.17632/5xfzcb4kn8.1 and https://doi.org/10.17632/8bhhk7c5n9.1). COVID-19 scRNA-seq datasets are publicly available from their original publications. All datasets used in this study are listed in Table 1.

### Authors’ contributions

A.B. and A.Z. conceptualized the study. A.B. implemented the method in code, analyzed data, and prepared the first draft of the manuscript. A.Z. supervised all research. All authors reviewed the manuscript.

---

**Figure 2—figure supplement 1.**
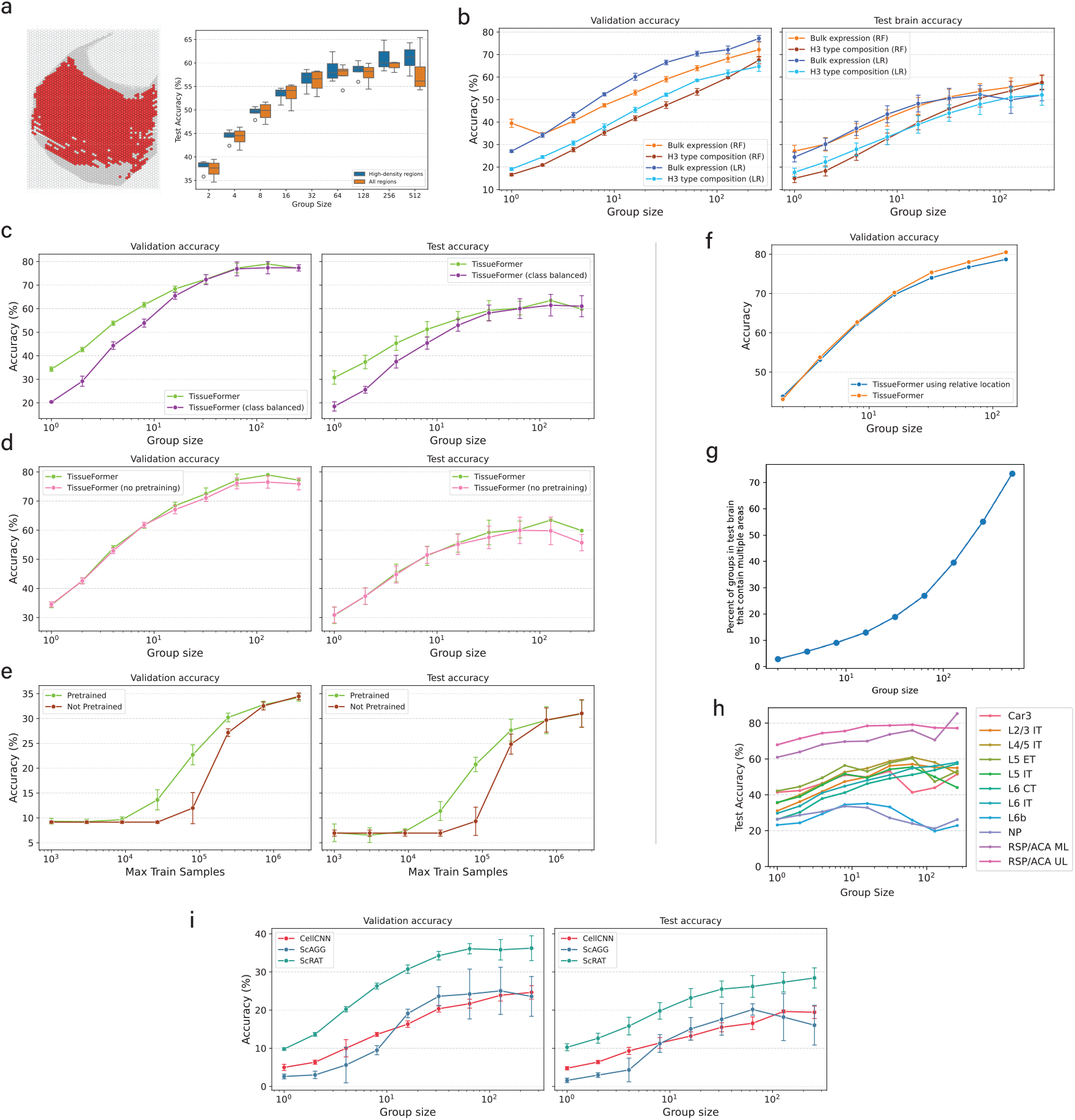
**a)** The drop in test brain accuracy relative to validation accuracy is only partially due to differences in sampling density. We selected hexes in the test brain which had high density in the 3 training brains (example at left), specifically, hexes with at least as many cells as the group size. (right) Test accuracy on data in those hexes was marginally higher than for all points, yet still much below validation accuracy. **b)** Performance of random forests (RF) compared against logistic regression (LR) given identical inputs. **c)** Performance of a class-balanced model which controls for some areas having more cells in training brains with a modified objective that discounts samples from frequent areas. **d)** Effect of initializing TissueFormer with a pretrained Murine Geneformer as single-cell module vs. randomly initialized. **e)** Varying the number of train samples, comparing random initalizations to pretrained initializations, here for *N* = 1 group size. *N* = 1 was chosen to reduce ambiguity and ensure that the number of cells seen equaled the number of groups seen. **f)** Effect of giving TissueFormer the relative spatial location (relative to the center of mass) within each group of each cell. **g)** Number of groups in the test brain that contain an area boundary as a function of the group size. **h)** The accuracy curves of models trained while only ever able to observe groups containing a single cell type. Groups were constructed by selecting a single cell and choosing the nearest N cells of the same time. The high performance of some types, such as RSP/ACA, reflects the fact that these cells are localized in just one area of cortex (retrospenial/anterior cingulate areas). Slopes of these curves are shown in Fig. 2. **i)** Validation (left) and test (right) accuracy as a function of group size for three deep learning benchmarks: CellCnn, scAGG, and ScRAT. TissueFormer (shown in ***Figure 2***f-g) outperforms all three methods.

**Figure 2—figure supplement 2.**
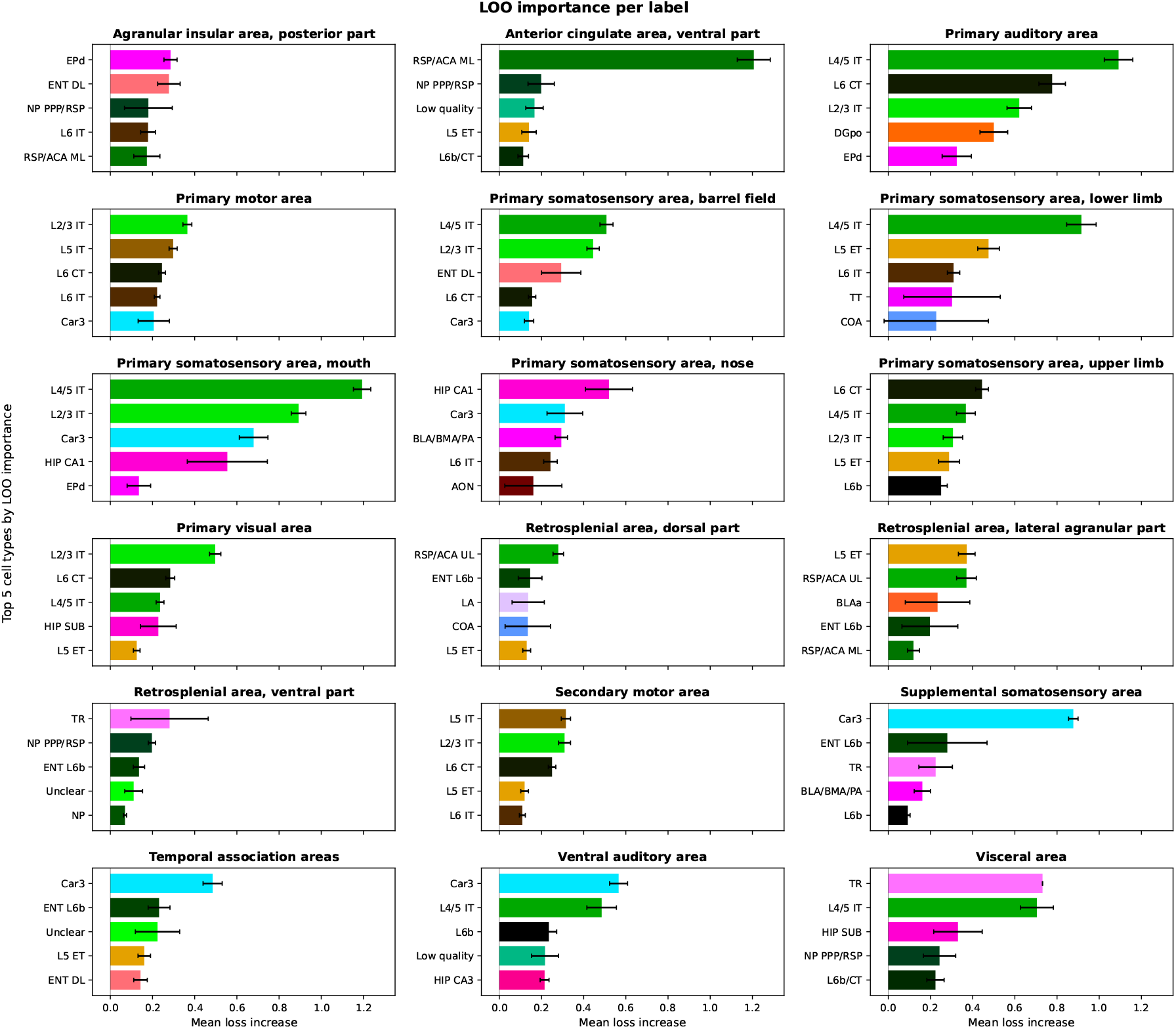
Leave-one-cell-type-out (LOO) analysis for the brain annotation task. Each panel shows the top cell types by LOO importance for a given predicted brain area. Importance is measured as the mean increase in cross-entropy loss when all cells of that type are removed from a group. L4/5 IT cells are consistently important across somatosensory areas, while L2/3 IT cells are most important for visual areas. Error bars represent standard error of the mean across groups.

**Figure 3—figure supplement 1.**
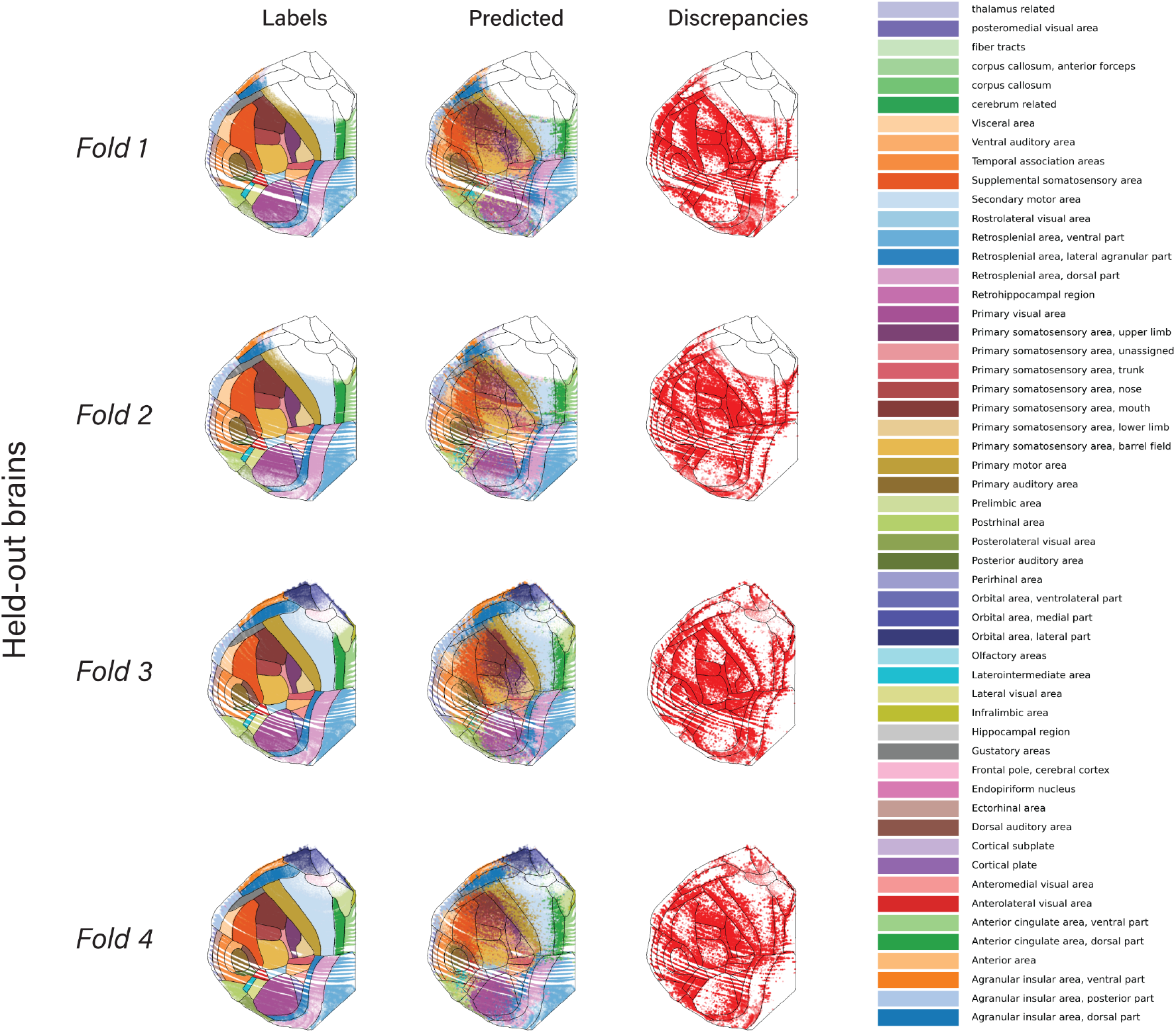
Across all 4 brains, we show the CCF labels on single cells (left), the predicted labels (middle) and the location of mismatches between the two. We call these discrepancies rather than errors as it is not clear which is closer to ground truth. Notable systematic differences include a gross shift of the Primary Visual Area (magenta), and the somatosensory-motor boundary. In both cases, a shift appears in the discrepancy map as a large amount of discrepancies on one side of an area but very few on the other side, and a large discontinuity at the CCF boundary.

**Figure 3—figure supplement 2.**
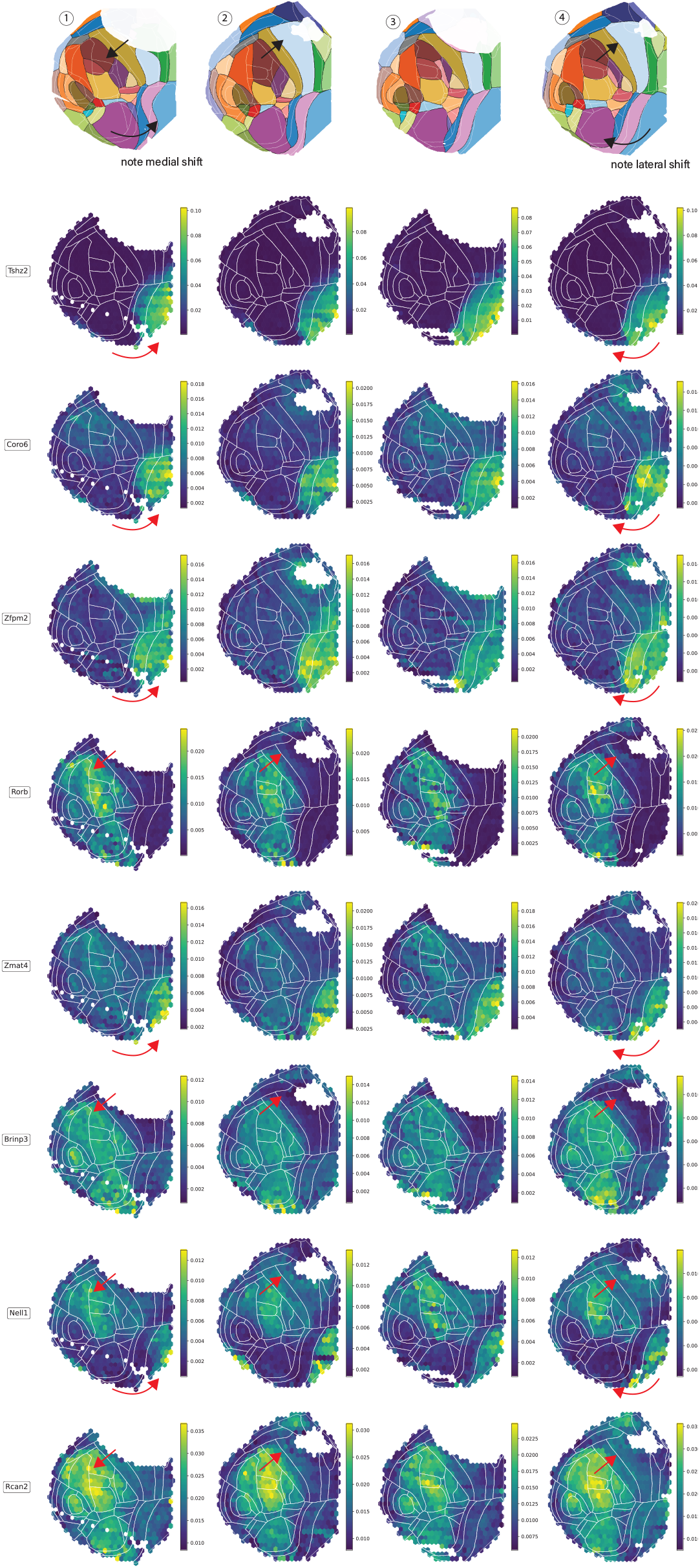
Normalized spatial density of select genes in each animal. Arrows highlight the animal-specific shifts of regional anatomy relative to the CCF (white boundaries), and how these are consistent between genes and with the predictions of Tissueformer (top row, reproduced from ***Figure 3***. The top three rows (Tshz2, Coro6, and Zfmp2) recapitulate the medial shift in the boundary to the retrospenial areas in Brain 1 and the lateral shift in this boundary in Brain 4. The bottom 5 rows recapitulate the shift in the somatosensory/motor boundary, the same as analyzed in cell type distributions in Figure 3. To construct normalized gene densities, we first normalized mRNA transcription levels within each cell. We then plot the average normalized transcription within each hex. Normalization was necessary to reduce confounds of unequal spatial sampling.

